# Multiplexed enrichment and tracking of lineages with CloneSweeper

**DOI:** 10.64898/2026.01.30.700779

**Authors:** Robert J. Vander Velde, Raymond W.S. Ng, Christopher Coté, Sydney M. Shaffer

## Abstract

A fundamental challenge in studying therapy resistance is understanding whether it results from pre-existing cellular states (“priming”) or drug-induced changes (“adaptation”). While lineage barcoding enables retrospective analysis of cells before and after treatment, current methods struggle to efficiently capture rare lineages in single-cell RNA sequencing (scRNA-seq) or isolate multiple specific lineages simultaneously for functional study. To overcome these limitations, we developed CloneSweeper, a multiplexed lineage tracking platform that pools enrichment libraries to isolate or enrich multiple rare lineages. CloneSweeper utilizes a dual-function barcode expressed as both a Cas9 gRNA for live-cell sorting and a 3’ UTR transcript for high-recovery detection in 10x Genomics scRNA-seq. We applied CloneSweeper to a model of BRAF V600E melanoma, where we identified that resistance to targeted therapy emerges from a polyclonal population of rare, pre-existing lineages. By simultaneously targeting and enriching 21 distinct rare lineages prior to treatment, we defined a heritable, primed state characterized by de-differentiation and elevated mesenchymal markers. We demonstrate that these primed cells are not quiescent but instead exhibit upregulated inflammatory and stress response signaling, specifically via the AP-1 and NF-κB1 pathways. CloneSweeper thus provides a robust framework for dissecting the molecular mechanisms of rare biological phenomena through simultaneous, multiplexed lineage isolation.

## Introduction

Cellular plasticity fuels adaptation in diverse biological contexts, ranging from developmental trajectories to the evolution of drug resistance in cancer. This plasticity confounds biological studies because observing cells only after they survive a stressor makes it difficult to distinguish between pre-existing features that conferred survival (“priming”) and transcriptional changes induced by the stressor itself (“adaptation”)^1,2,3,4,5,6^. In cancer, for example, distinguishing whether a surviving cell possessed intrinsic resistance or adapted during therapy is largely impossible without retrospective analysis (Figure 1A). Simply studying cells prior to treatment remains insufficient when the pre-perturbation population is heterogeneous^7^, especially when the population of interest is rare, such as priming to therapy resistance (Figure 1A)^2,8^.

**Figure 1:**
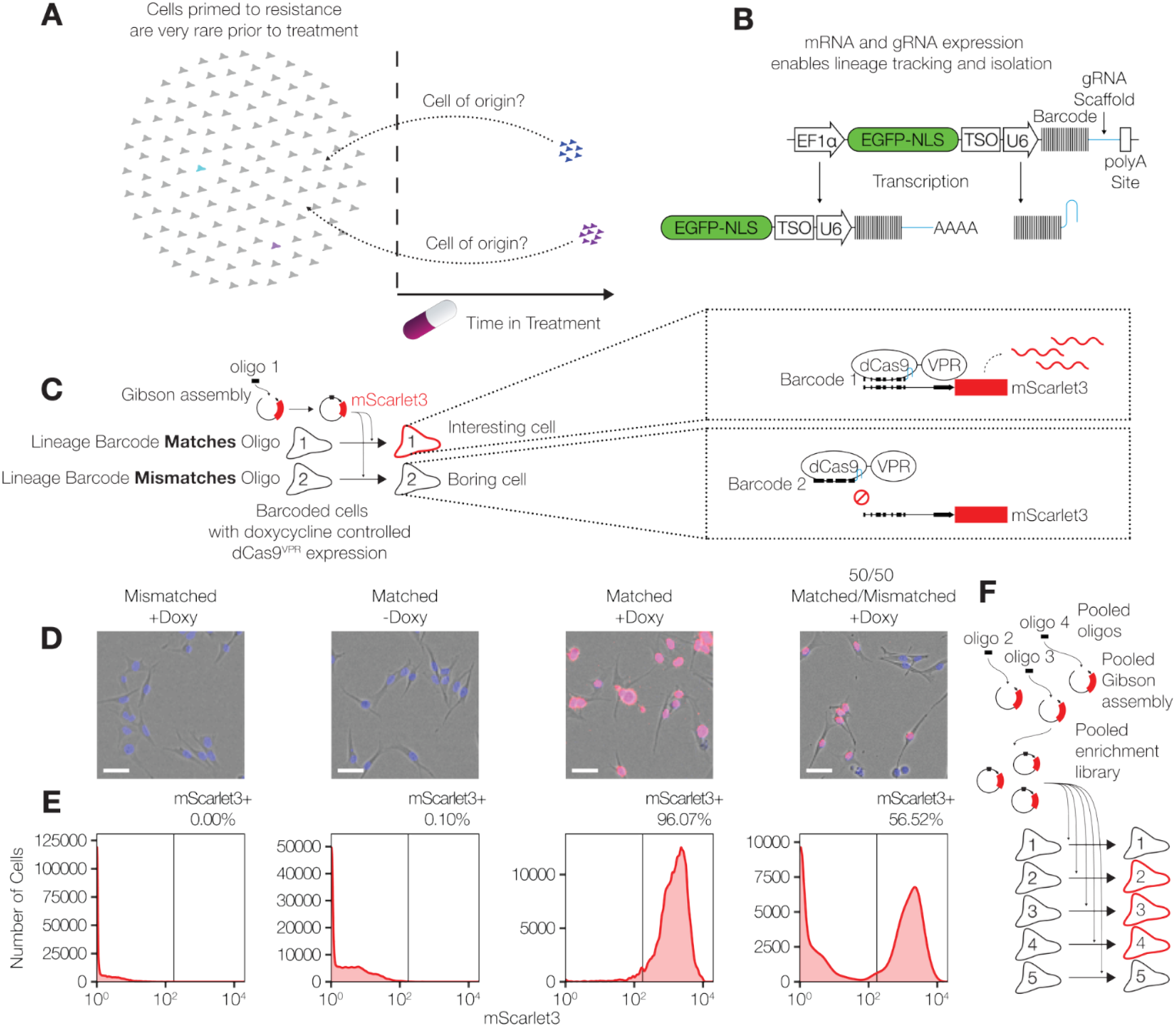
CloneSweeper enables lineage tracking and isolation via dual-function barcodes. **A**. Schematic illustrating the challenge of distinguishing pre-existing priming from therapy-induced adaptation in rare subpopulations. **B**. Design of the CloneSweeper lineage barcode. The barcode sequence is expressed as both a poly-adenylated transcript (for scRNA-seq detection) and a Cas9 gRNA (for live-cell isolation). **C**. Strategy for targeted isolation. Gibson assembly is used to clone identifying barcode sequences into a reporter plasmid. In the presence of the specific gRNA barcode, dCas9-VPR binds the reporter, driving mScarlet3 expression. **D**. Representative Incucyte images of mScarlet3 activation 32h post-doxycycline induction. EGFP (lineage barcode marker) is pseudocolored blue, and mScarlet3 (target reporter) is red. Scale bars are 56.4 µm. **E**. Flow cytometry analysis of mScarlet3 expression 48h post-induction, demonstrating high specificity for the matched lineage. **F**. Schematic of the pooled enrichment strategy. Multiple reporter plasmids are pooled to simultaneously isolate diverse lineages from a heterogeneous population.

Lineage barcoding addresses this challenge by using heritable barcodes (usually integrated into DNA^9,10,11,12,13,14^) to identify closely related cells. This approach enables retrospective analysis by determining which cells in a pre-perturbation state or control state relate to those that respond to a perturbation in a specific way, such as surviving in therapy. Recent advances in lineage barcoding have significantly expanded our capacity to perform these retrospective analyses.

Early techniques, such as ClonTracer^9^, integrated lineage barcodes into DNA. By assuming that DNA reads correlate with lineage population size, researchers can independently track the growth dynamics of lineages with and without perturbation. Updated technologies, such as CellTag^10^ and LARRY^11^, allow researchers to detect lineages in single-cell RNA sequencing (scRNA-seq) data by expressing their barcodes as RNA. By reading the transcriptomes prior to alterations or treatments, these methods add to our knowledge of pre-treatment gene expression. Other techniques, such as REWIND^12^, express long barcodes to detect via RNA FISH^15^, a method later adapted for scRNA-seq^16,17,18,19^. With the advent of combined single-cell ATAC-Seq and RNA sequencing (multiomics), these barcoding technologies now support a deeper understanding of how a cell’s lineage, gene expression, and chromatin accessibility interact^20^.

However, these techniques struggle when the phenotype of interest is rare prior to treatment. This limitation is particularly evident in the context of priming to therapy resistance, as standard sampling often fails to capture these scarce cells (Figure 1A). To overcome this, methods like ClonMapper^13,21^ and CaTCH^14^ functionalize the lineage barcode, transforming it from a passive label into a tool for physical isolation. By expressing barcodes as Cas9 guide RNAs (gRNAs) or linking them to marker genes, these systems allow researchers to selectively activate fluorescence or drug resistance in specific clones. This enables the physical enrichment of rare lineages from the bulk population, making them accessible for analysis. Yet, until now, these methods have only isolated single lineages at a time.

Current lineage tracing tools face three key limitations. First, scRNA-seq sequencing can capture barcode sequences inefficiently, often requiring deep sequencing or side reactions to recover the barcode of interest. Second, because lineages of interest are often rare in the bulk population, capturing sufficient representation for statistical analyses requires sequencing a prohibitively large number of cells. Third, existing methods for isolating specific lineages only target one clone at a time, limiting their utility for studying polyclonal phenomena and reducing experimental efficiency.

To overcome these limitations, we developed CloneSweeper, a lineage tracing platform that addresses these three challenges. CloneSweeper tracks clones through time, provides high recovery in 10x Genomics scRNA-seq, and enables polyclonal live-cell recovery. We validated CloneSweeper in a model of targeted therapy resistance in BRAF V600E melanoma, where cells primed for resistance constitute an extremely rare subpopulation prior to treatment^2,16,17^.

## Results

### Generating a Lineage Barcoding Library Capable of Isolating or Enriching Rare Lineages

To overcome the rarity of primed cells and track lineages before and after treatment, we developed CloneSweeper. This system uses a dual-function lineage barcode expressed as both a gRNA for a Cas protein and as a part of the 3’ UTR of EGFP (Figure 1B). This design achieves two important functions for CloneSweeper. First, for scRNA-seq, we positioned the gRNA within 225 bases of the polyadenylation signal and adjacent to a 10x Genomics Template Switch Oligo (TSO) site. This design ensures that the lineage barcode is included in the 3’ sequence captured for sequencing. Second, for live-cell isolation, the gRNA directs a catalytically dead Cas9-VPR fusion protein to a specific target sequence. When we add a “reporter” plasmid containing a target sequence matching the lineage barcode, the Cas9-VPR/gRNA complex binds the promoter activating expression of a fluorescent protein or other target sequence. This enables FACS-based enrichment of cells from any lineage of interest. We hypothesized that these design characteristics would enable both efficient lineage tracking and lineage isolation to study priming for drug resistance.

### Validating CloneSweeper’s ability to activate a marker in cells with a specific barcode

We tested CloneSweeper’s specificity using single-cell derived subclones of lineage barcoded WM989 melanoma cells with integrated doxycycline inducible Cas9-VPR. After identifying the lineage barcodes of these subclones via Sanger sequencing, we used Gibson assembly to clone a 20bp sequence matching specific barcodes upstream of a minimal CMV promoter and mScarlet3. We designed this sequence to match one lineage barcoded (matched reporter in Figure 1D,E) but not the other lineage barcode (mismatched reporter in Figure 1D,E).

To test the specificity of the marker activation, we transduced these two subclones with their respective mScarlet3 reporter plasmids. This resulted in a high degree of separation in mScarlet3 fluorescence between the matched and mismatched lineages, demonstrating that CloneSweeper is highly specific to the integrated lineage barcode (Figure 1D,E). These data also show that the system remains tightly regulated, as the subclone with a matched barcode exhibited minimal fluorescent leakage in the absence of doxycycline (Figures 1D,E).

CloneSweeper can be scaled to isolate or enrich multiple rare subclones simultaneously. This is achieved through the pooled cloning of recovery plasmids, which provides a significant advantage over previous approaches that target only a single lineage at a time (Figure 1F). We utilized this novel, multiplexed approach for all subsequent downstream experiments.

### Identifying lineages primed to resistance

We applied CloneSweeper to study the evolution of resistance to dabrafenib and trametinib combination therapy in BRAF V600E positive melanoma cells. We first transduced doxycycline inducible Cas9-VPR containing WM989 cells with the CloneSweeper lineage barcode library, followed by sorting for EGFP positive cells (selecting for cells transduced with a lineage barcode). After approximately 8.5 doublings, we plated cells into replicates and then treated them with dabrafenib and trametinib the next day. At this point we collected a cell pellet for analysis of lineage population sizes prior to treatment. We also cryopreserved a live subset of cells before treatment to use for enrichment at a later time (Figure 2A). After six weeks of treatment, we harvested the resistant cells from each replicate. We simultaneously plated an additional pool of cells after sorting for EGFP positive cells for a semi-independent experiment. Results of this experiment are shown in Figure S1.

**Figure 2:**
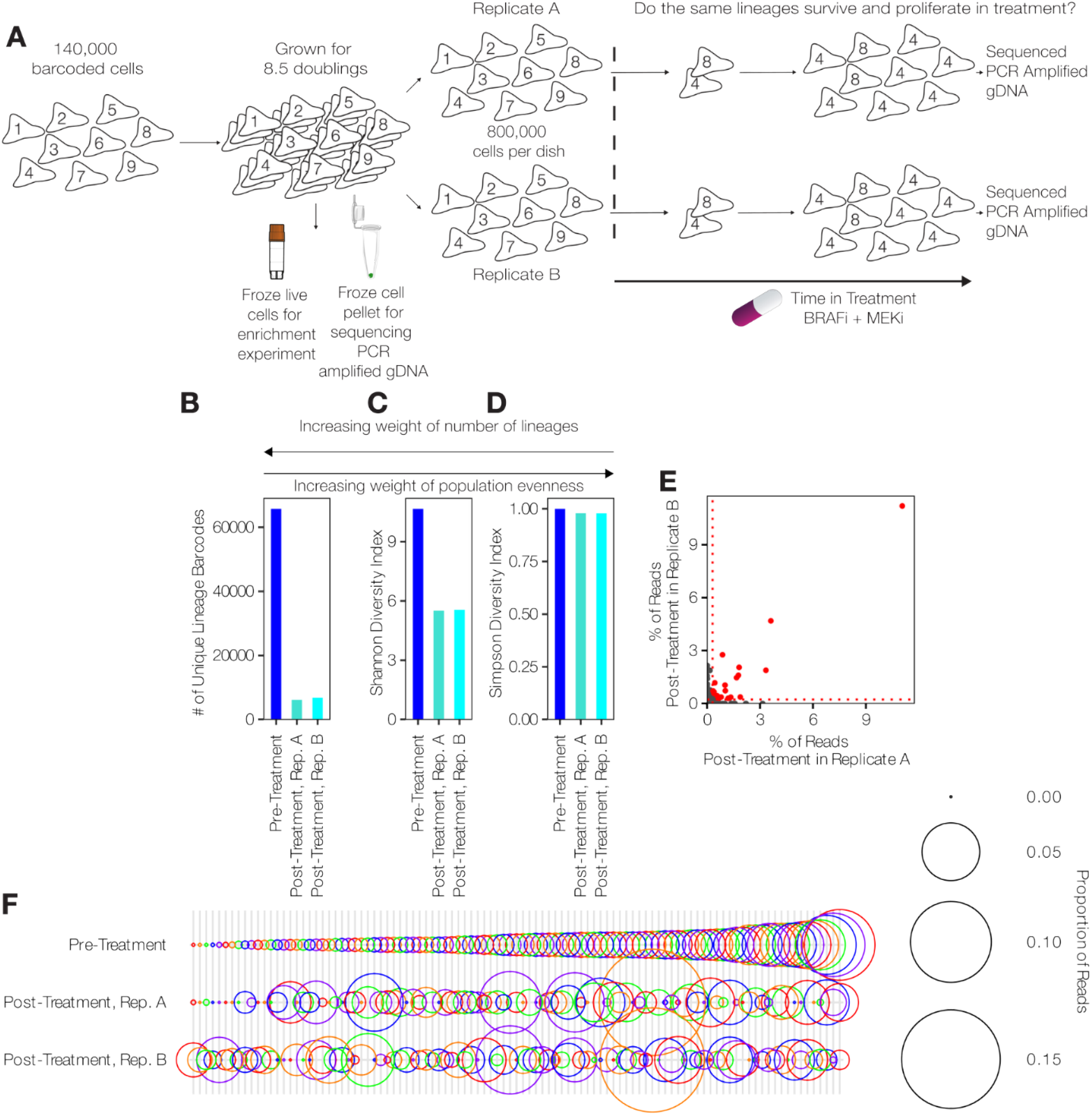
Longitudinal lineage tracking identifies heritable resistance in BRAF V600E melanoma. **A**. Schematic of the experimental workflow. A barcoded population was expanded (∼8.5 doublings) and split into a pre-treatment baseline (sequenced and cryopreserved) and two replicates treated with dabrafenib and trametinib. **B**. Number of unique lineage barcodes detected in pre- and post-treatment samples. **C**. Shannon diversity index demonstrating population bottlenecks. **D**. Simpson diversity index indicating the maintenance of polyclonal diversity. (B-D) Metrics include only barcodes with >= 2 reads. **E**. Correlation of lineage abundance between post-treatment replicates. The red dotted line represents the 200-cell threshold derived from spike-in controls. Red dots indicate the 21 lineages identified as “primed” (present >200 cells in both replicates). **F**. Bubble plot displaying the frequency of surviving lineages relative to their pre-treatment abundance. Colors cycle across adjacent lineages for visual distinction.

We identified lineages that survived therapy by isolating genomic DNA and PCR amplifying the CloneSweeper lineage barcodes and sequencing. We quantified lineage abundance by associating the amplified read counts with the specific cell count for each lineage. We then compared the diversity metrics of the cell populations before therapy to those that survived and became resistant to therapy. Of the metrics used here, the Shannon index is considered highly sensitive to the total number of unique lineages, while the Simpson index relies heavily on the diversity of the most abundant clones^22^. We found that the total number of unique lineage barcodes and the Shannon diversity index decline precipitously in the posttreatment samples compared to the pre-treatment condition (Figure 2B,C). The dramatic reduction in unique barcodes and Shannon index reflects the substantial loss of lineages during treatment, presumably in drug susceptible lineages. In contrast, the Simpson index showed a much smaller reduction (Figure 2D). The stability of the Simpson index indicates that the surviving population remains highly diverse and polyclonal rather than being dominated by a single expanded clone. This result highlights a critical limitation in current single lineage isolation methods. Tools that isolate only one clone would fail to capture the heterogeneity of this resistant state. To address this, we prioritized a strategy capable of identifying and enriching multiple lineages simultaneously for future enrichment experiments.

We next asked whether we detected the same lineages across our replicates to further assess if resistance arises stochastically or deterministically in specific lineages^9,23^. We found that surviving lineages between replicates were sufficiently correlated to suggest that a heritable primed state exists (Figure 2E). One challenge in analyzing barcode data from genomic DNA is that we generally do not know how barcode reads relate to cell numbers. To roughly correlate reads with cell numbers, we included a benchmarking “ladder” with cells of known barcodes at different numbers (20, 100 and 200) and we added these cells to our genomic DNA isolation.

Since these cells are amplified with the library, we can use these to estimate the abundance of each barcode across samples. Using the mean read count of the 200-cell ladder rung as a lower limit threshold (Figure 2E, red dotted line), we identified 21 lineages that exceeded this estimated size in both replicates, further confirming the heritability of the resistance phenotype. To rigorously test for heritability, we calculated the concordance of resistance at the 200 cell cutoff. We found that if a lineage passed the abundance threshold in one replicate, it was overwhelmingly likely to pass it in the other (odds ratio 920.47). Importantly, we observed minimal concordance between initial barcode representation prior to treatment, indicating that the initial growth rate is not the primary determinant of resistance (Figure 2F). We therefore selected these 21 lineages for simultaneous enrichment from the naive population using a custom, pooled CloneSweeper library.

### Enrichment of specified lineages

We next validated the multiplexing capacity of CloneSweeper by targeting the 21 different lineages identified in our resistance experiment (Figure 2E). We constructed a pooled enrichment “mini library” by using Gibson assembly to insert oligos matching these barcodes into the same mScarlet3 reporter backbone we validated earlier (Figure 1F). Sequencing of this plasmid library confirmed that we successfully included all 21 of the lineage barcode sequences in this library (Figure S2B), indicating that we successfully cloned a bespoke, pooled enrichment mini-library based on lineages of interest.

We then thawed the original pre-treatment cells and transduced them with this enrichment library for isolation of the corresponding lineages (Figure 3A). With this experimental design, we expect cells corresponding to the 21 lineage barcodes in the library to express the mScarlet3 fluorescent protein. We used FACS to isolate these mScarlet3 and EGFP positive cells (found at 0.1% to 0.53% of the EGFP positive population). Because the rare lineages provided insufficient material for the optimal use of 10x scRNA-seq reagents, we reached the optimal cell count by pooling the isolated mScarlet+ cells with a population of mScartlet-cells. This approach maintained our enriched lineages of interest while simultaneously providing an internal control of non-primed cells within the same reaction, thereby eliminating batch effects when comparing transcriptional states. We extracted CloneSweeper lineage barcodes from the raw sequencing reads and integrated this identity information directly into our scRNA-seq analysis pipeline. After filtering for standard scRNA-seq quality metrics (see methods), we obtained very high detection of CloneSweeper lineage barcodes (Figure S2C-E). We confidently assigned 75.68% of cells post-Seurat filtering to a single lineage barcode (see Methods).

**Figure 3:**
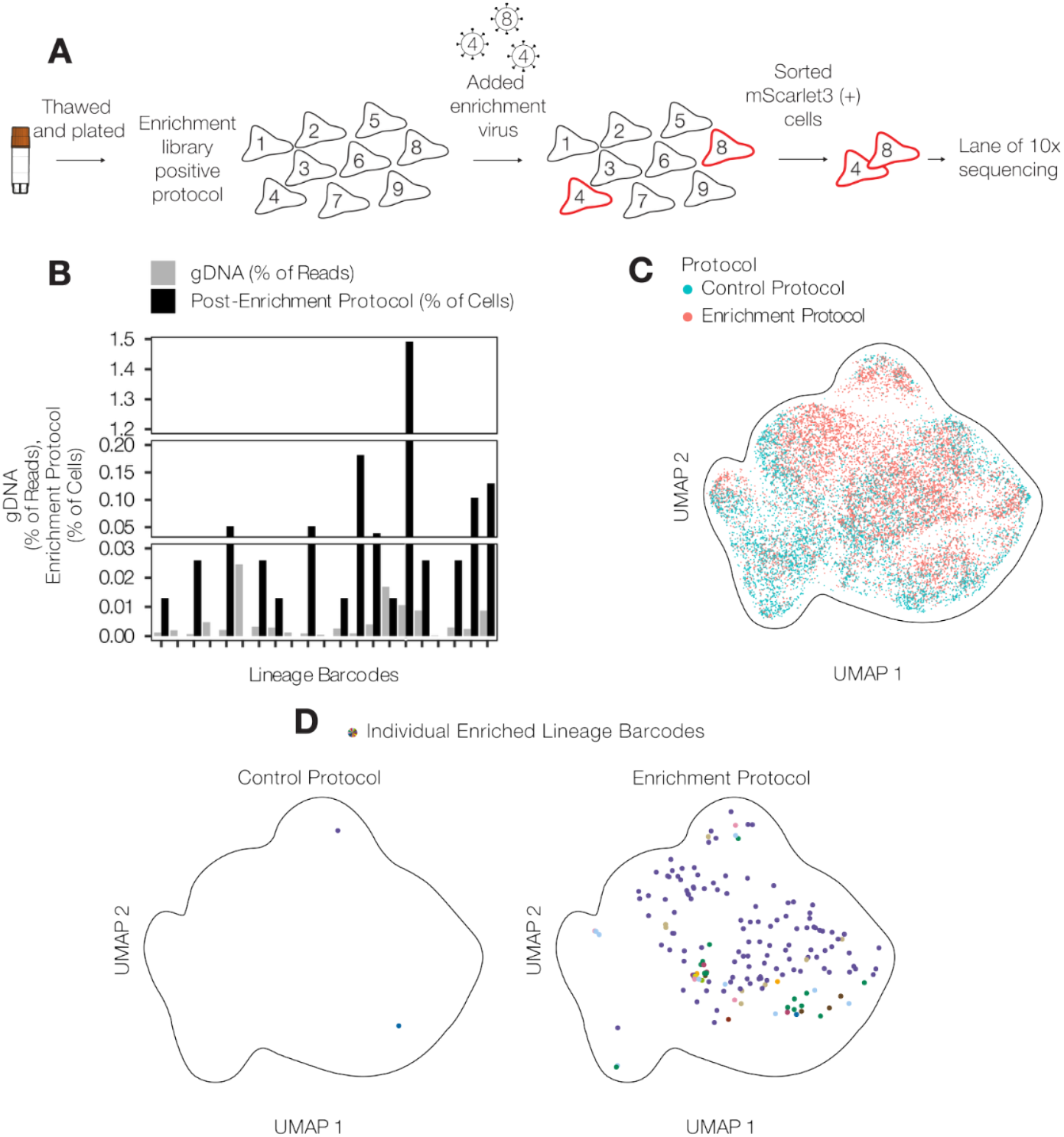
Multiplexed enrichment recovers rare primed lineages for scRNA-seq. **A**. Schematic of the experimental workflow for enrichment. Live pre-treatment cells were thawed and transduced with the pooled “mini-library” targeting the 21 identified resistant lineages, followed by FACS isolation of mScarlet3+ cells. **B**. Comparison of lineage abundance. Black bars show the frequency of targeted lineages in the bulk pre-treatment gDNA; colored bars show the frequency of the same lineages in the enriched scRNA-seq dataset (expressed as % of cells). **C**. UMAP projection of scRNA-seq data (cell cycle regressed) comparing the cells from the control protocol (blue) to those from the enrichment protocol (pink). **D**. Location of the 21 targeted lineages within the UMAP projection. Note the sparse detection from the control protocol (left) versus the dense recovery from the enriched protocol (right). Outline is aligned to correspond exactly to the outline in C.

To test whether physical enrichment is required to capture rare primed cells, we performed a side-by-side comparison between our enriched sample and a control population that was not transduced with the enrichment library (Figure S3A). We used this design to determine the baseline frequency of the primed lineages that would be captured by chance while also measuring the transcriptional effects of transduction. Within the enriched sample, we detected 15 of the 21 targeted lineage barcodes. Compared to our bulk genomic DNA estimates, 14 of these 15 lineages were substantially more abundant in the scRNA-seq from the enrichment sample (Figure 3B). We visualized the global transcriptional landscape of both the enriched and control populations using UMAPs (Figure 3C, S3A). The enriched sample displayed a dense distribution of cells from multiple targeted lineages (Figure 3D,S3B). In contrast, the control sample contained only two cells with the targeted lineages, representing an >80 fold increase in the number of enriched cells. This drastic difference demonstrates that CloneSweeper effectively retrieves the diverse array of rare lineages that drive resistance which standard scRNA-seq fails to capture.

### Investigation of cells primed to resistance

We next characterized the transcriptional identity of these enriched cells to understand how they survive therapy. Differential expression analysis revealed 317 genes with significantly higher or lower expression in enriched cells primed for resistance (Figure 4A, S3C,D). Broadly, the enriched cells were in a de-differentiated state compared to the rest of the population. Specifically, these cells displayed significantly lower expression of markers of melanocyte differentiation *DCT* and *PMEL* (though *TYR* was not significantly different) (Figure 4B). Concurrently, they exhibited significantly higher expression of the mesenchymal markers *CD44* and *CD59* (Figure 4C). This phenotype mirrors dedifferentiated states previously described in the literature^24,16,25,26^, where the loss of lineage identity coupled with gained mesenchymal features marks a plastic state capable of surviving drug pressure.

**Figure 4:**
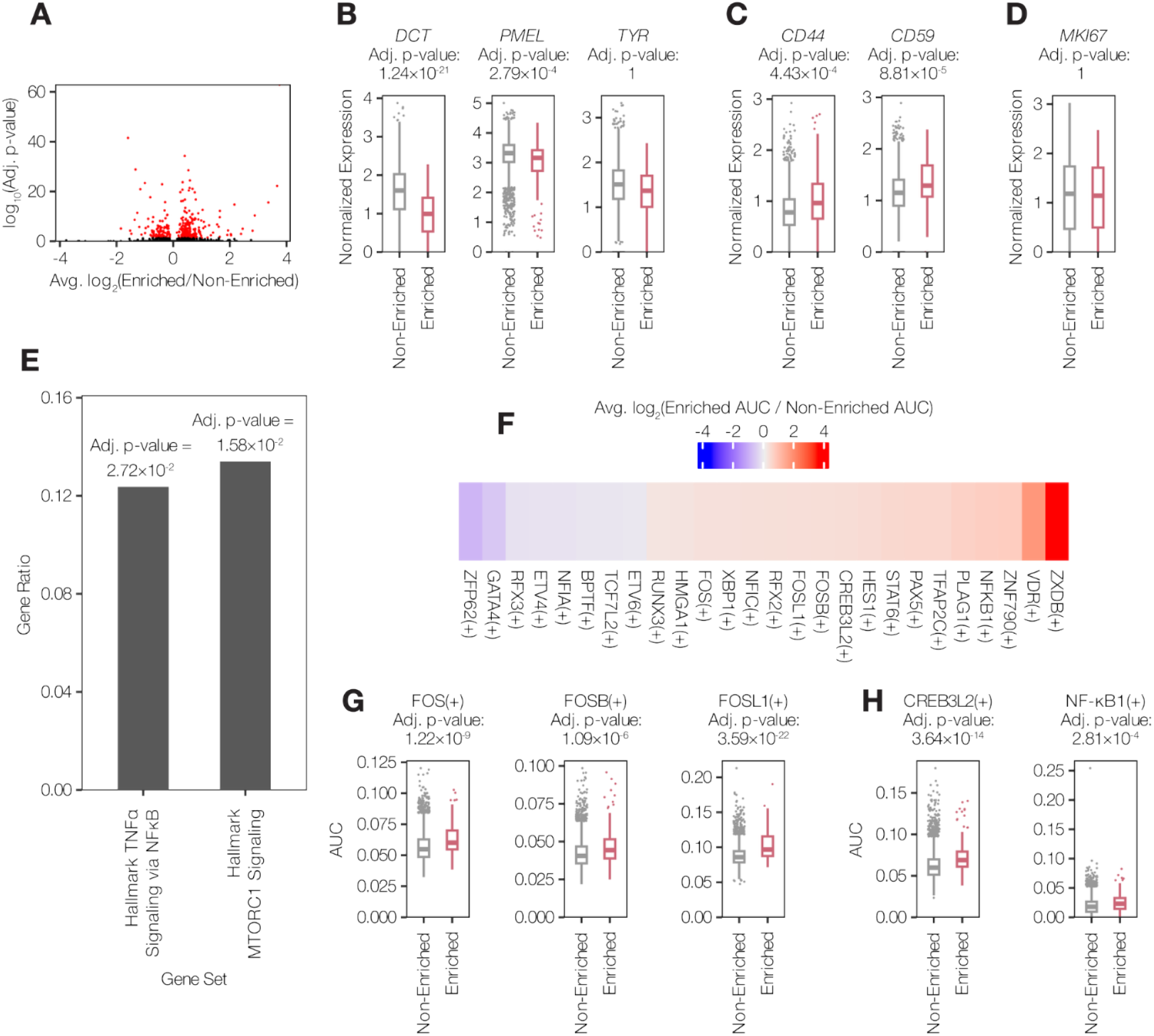
Primed lineages exhibit a de-differentiated, stress-responsive transcriptional state prior to treatment. **A**. Volcano plot showing differential gene expression between enriched (primed) and non-enriched cells within the library-positive sample. P-values determined by Wilcoxon rank-sum test with Bonferroni correction. Only genes detected (UMIs ≥ 1) in 5% of cells in both groups are shown. **B-D**. Normalized expression of (B) melanocyte differentiation markers (*DCT, PMEL, TYR*), (C) mesenchymal markers (*CD44, CD59*), and (D) the proliferation marker MKI67. Boxplots show median and interquartile ranges, whiskers show 1.5× the IQR. P-values are determined by the Wilcoxon rank-sum test with Bonferroni correction. **E**. GSEA showing significantly upregulated Hallmark pathways in primed cells (Adjusted p-value < 0.05). **F**. Heatmap of differential transcription factor activity inferred by SCENIC (Area Under the Curve, AUC). Colors represent the log_2_ fold change between enriched and non-enriched cells. **G-H**. Comparison of SCENIC AUC distributions for (G) AP-1 subunits (FOS, FOSB, FOSL1) and (H) stress response factors (CREB3L2, NF-κB1). P-values are determined by the Wilcoxon rank-sum test with Bonferroni correction. Boxplots show median and interquartile ranges, whiskers show 1.5× the IQR.

A classic model of therapy resistance suggests that survival is driven by a slow-cycling ‘persister’ state^8,27^. However, in many of these studies, it remains unclear whether these persisters are intrinsically slow cycling prior to treatment^8^. We explicitly tested for pre-existing differences in cell cycle progression by characterizing the transcriptional state of these lineages before drug exposure. Crucially, we found that primed cells expressed the cell cycle marker MKI67 at levels indistinguishable from the non-primed population (Figure 4D). This finding demonstrates that these lineages were not slow-cycling prior to treatment. Consequently, the reduced proliferation observed in resistant colonies is not a mechanism of initial survival, but rather a tolerance phenotype induced by the therapy itself.

### Inferred Regulatory Networks in Primed Cells

We next performed Gene Set Enrichment Analysis (GSEA) to identify the signaling cascades driving this primed state. Enrichment analysis using lists of significant genes (adjusted p-value < 0.05) revealed enrichment for hallmarks of TNFα signalling via NFκB and mTORC1 signalling in primed cells prior to treatment (Figure 4E). However, we found no Hallmark pathways significantly enriched in non-primed cells compared to primed cells. Upregulated hallmarks in primed cells suggest a possible role for inflammation^28^ and metabolic heterogeneity^29^ in priming to resistance.

To determine the upstream drivers of these programs, we performed SCENIC analysis to infer transcription factor activity. This analysis revealed distinct regulatory networks specific to the primed cells (Figure 4F). We inferred increased activity of FOS regulons (Figure 4G) in primed cells. This suggests elevated AP1 signaling prior to treatment in this population, which aligns with previous studies implicating stress response pathways in adaptive resistance^30,26^. We also inferred increased activity of CREB3L2 and NF-κB1 (Figure 4H) in primed cells. CREB and NF-κB1 regulation have been implicated in resistance to BRAF inhibitors previously^31,32^. Collectively, through enrichment of this primed population and scRNA-seq, we comprehensively define this heritable primed state that enables melanoma cells to escape targeted therapy.

## Discussion

Here we developed CloneSweeper, a versatile lineage tracing platform for simultaneous tracking, scRNA-seq, and enrichment of multiple rare lineages. Our platform achieves high barcode recovery in scRNA-seq without the need for excessively deep sequencing or a PCR side-reaction. It also has the ability to isolate live cells with lineages of interest in a pooled assay. We demonstrated these functionalities in a BRAF V600E melanoma model treated with targeted therapy. In this system, CloneSweeper’s multiplexed enrichment is essential as resistance is a polyclonal phenomenon. By targeting 21 clones simultaneously, we enriched the primed cells >80-fold over their baseline frequency, enabling robust characterization of a population that would be missed by standard scRNA-seq.

Our analysis of primed melanoma cells revealed a de-differentiated transcriptional state characterized by reduced melanocyte markers and elevated mesenchymal markers. This phenotype aligns with previous descriptions of plastic, therapy-resistant states in melanoma and other cancers^33,16,34,35^. Importantly, by analyzing cells before drug exposure, we distinguished true priming factors from drug-induced responses. Primed cells showed no difference in proliferation markers prior to treatment, indicating that resistance does not require pre-existing quiescence. Instead, these cells exhibited elevated AP-1, NFκB, and mTORC1 signaling, suggesting that inflammatory and metabolic programs contribute to the primed state. CloneSweeper allows for the enrichment of live-cells, in this case for straightforward 10x scRNA-seq. This approach enables sequencing at a higher depth per cell, by eliminating the need to sequence a huge number of cells (as is possible with combinatorial based scRNA-seq).

CloneSweeper offers distinct methodological advantages over existing lineage tracing technologies. Relative to the REWIND or FateMap technique^12,16,17,18,19^, CloneSweeper barcodes can be identified directly from 10x scRNA-seq without requiring a side PCR reaction or sequencing of large constructs^16,17^ due to CloneSweeper barcodes’ close proximity to a polyA tail. CloneSweeper is similar to ClonMapper^13,21^ in that it expresses lineage barcodes as gRNAs. We expanded on this established utility by implementing pooled “enrichment libraries” to target multiple lineages simultaneously. This multiplexing capacity allows us to efficiently sequence multiple clones in parallel with minimal expansion, thereby preserving the transcriptional state of the cells.

We also distinguish our approach from systems like CaTCH. CaTCH elegantly uses the lineage barcode as a target site within a promoter rather than expressing it as a gRNA^14^. This design ensures the barcode remains functionally inert until targeted, thereby preventing potential off-target effects that can arise when lineage barcodes are constitutively expressed as gRNAs. While the CaTCH design prioritizes this safety, CloneSweeper adopts an alternative architecture that prioritizes modularity and direct transcriptomic detection. Because we express the barcode as a transcript, we can identify it immediately in scRNA-seq, while CaTCH cannot. This flexibility also permits the exchange of marker genes and plasmid backbones without altering the primary library, enabling diverse functional assays from a single resistance experiment.

CloneSweeper also has limitations that warrant consideration. First, expressing lineage barcodes as gRNAs renders CloneSweeper relatively incompatible with other Cas9-based assays, although this could be avoided in the future using Cas9 orthologs or Cas12 in the recovery system. Second, we observe some lineage barcodes that are enriched more efficiently than others (Figure 3B,D), which could be problematic for certain studies. Third, similar to other barcoding technologies^9,10,11,16,36,13,14^, CloneSweeper relies on medium to long-term heritability of cell-states prior to treatment to allow enough cell growth to detect rare cell states. This disadvantage can be circumvented with technologies that track cell-states in ancestor cells as opposed to sibling cells^37^, though these techniques have their own unique limitations.

In designing CloneSweeper experiments, there are important considerations that are critical to obtaining interpretable results. First, the phenomenon of interest must be sufficiently heritable through the same number of divisions as the experiment. This number of divisions is important because the clones need to expand enough to make replicates for the experiment and for cryopreservation. Second, the number of barcodes in the enrichment library should be carefully selected. Here, we enriched for 21 barcodes, but this required a very high multiplicity of infection to ensure that each cell received a reasonable representation of each of the recovery plasmids. Fewer barcodes enriched might be advisable to decrease potential off-target effects and ensure high recovery of the lineages of interest. Furthermore, tighter gating of mScarlet3 positive cells might ensure higher enrichment of cells. We therefore believe that future CloneSweeper experiments can be further optimized based on our observations here.

In conclusion, we have developed a robust system for simultaneously and efficiently enriching multiple rare subclones of interest for scRNA-seq. We tested this system in the context of the evolution of resistance to targeted therapies in melanoma. We foresee this technology having a broad impact across multiple contexts, including resistance to other treatments, cancer initiation and developmental biology.

## Methods

### Cell culture

We obtained WM989 cells from the A6-G3 subclone derived earlier ^2^. We cultured WM989 cells in Tu2%, prepared as reported earlier^2,38,36^. We passaged and collected WM989 cells using 0.05% trypsin (Gibco 25300054). Dabrafenib and trametinib resistant cells were collected using 0.25% trypsin (Gibco 25200056). We cultured HEK293T and LentiX in DMEM/F-12, GlutaMAX(TM) (Gibco 10565) with 10% FBS (Cytiva HyClone SH3039603) and 1% Penn/Strep (Gibco 15140122). We passaged them using 0.05% trypsin.

### Generation of lineage barcoding library (CloneSweeper)

We derived the backbone of CloneSweeper from CROPseq-Guide-Puro^39^ (Addgene #86708). We replaced the puroR with EGFP-6xMYC-NLS from VB211026-1265ped (constructed by VectorBuilder). We used Phusion PCR (NEB M0531S) followed by In-Fusion (Takara Bio 638909) cloning to replace the PuroR. We inserted a TSO site using the same techniques with the oligos: GACCCAGAGAGGGCCTATTTCC and GGCCCTCTCTGGGTCCCCATGTACTCTGCGTTGATACCACTGCTTCCCTCGGGGTTGGGAGGTGG. We then linearized the plasmid using BsmBI-v2 (NEB R0739S). We extracted linearized plasmid from agarose gel using a Qiagen gel extraction kit (28704); we eluted DNA in 30µl of molecular grade water heated to 65°C, which we applied to multiple columns. We then prepared three 50µl Gibson assembly (NEB E2611S) reactions, each with: 250ng of linearized plasmid, 971fmol of oligo and 25µl of Gibson (NEB E2611S) master mix. We used the following HPLC purified oligo mix as an insert: TGGAAAGGACGAAACACCG NSNW NSNW NSNW NSNW NSNW GTTTTAGAGCTAGAAATAGCAAGTTAAAATAAGGC. We also prepared a 50µl control reaction without oligo. We precipitated all 50µl reactions separately and resuspended them in 10µl water. We transformed Endura electrocompetent bacteria (Bioresearch Technologies 71003-036) with 2µl of concentrated DNA per 25µl bacteria. We performed twelve transformations and added these to 100µg/ml ampicillin LB broth for maxi-prep plasmid extraction (Invitrogen K210006). Plating 1:10,000 of recovery media from each transformation suggested a total complexity of 27,982,000 barcodes, after adjusting for the number of colonies produced in the negative control at a 1:1000 dilution.

We produced CloneSweeper lentivirus via transfection of HEK293T cells in four nearly confluent 10cm plates. We added 2ml of Opti-MEM to two tubes. We added 320µl of pH 7.0 1mg/ml PEI (Polyscience 23966-1) to one tube. We vortexed this tube and incubated the solution for 5min. We added 40µg of CloneSweeper library to another tube as well as 30µg of pPax2 and 20µg of VSVG. After combining the contents of the two tubes, we vortexed and incubated the solution for 15min. We then added 1ml of the combination dropwise to each plate. Approximately 6hrs later we replaced the media with 5ml Tu2%. We collected virus every 5-17 hrs. After six collections we filtered virus through a 0.45µm mesh (Millipore Sigma SE1M003M00) and aliquoted.

### Generation of WM989 cells with doxycycline inducible Cas9-VPR

We plated WM989 A6-G3 at 100,000 cells per well in a 12-well plate. We added 30µl of Opti-MEM (Gibco 31985062) to two tubes. We added 5µl of pH 7.0 1mg/ml PEI (Polyscience 23966-1) to one tube, we vortexed this tube and incubated solution for 5min. We added 1µg of Piggy-bac transposase plasmid (Systems Biosciences PB210PA-1) and 1µg of PB-TRE-Cas9-VPR (Addgene #63800) to the other tube. We mixed the tubes together and vortexed it, we then incubated the solution for 15min. We then added 60µl of this mix dropwise. We grew the transfected and untransfected (control) cells in the presence of 500µg/ml hygromycin B (Corning MT30-240-CR). This selection eliminated all control cells.

### Testing clone identification with CloneSweeper

We transduced PB-TRE-Cas9-VPR WM989 A6-G3 with CloneSweeper at 2.34% transduction efficiency. We then sorted EGFP positive cells into single wells of a 96 well plate using a FACSJazz sorter. Using early Incucyte scans, linked to later scans of colonies, we confirmed that colonies were derived from single cells. We extracted genomic DNA from these colonies using a QiaAmp DNA Mini Kit (Qiagen 51304). We used Phusion PCR (NEB M0531S) to amplify the CloneSweeper barcodes. We used Sanger sequencing to identify the CloneSweeper barcodes integrated into each subclone. VectorBuilder constructed VB230528-1280dkp which we used as a source of EGFP-6xMYC-NLS. We linearized this backbone with BsmBI-v2 (NEB R0739S). We gel extracted using the Qiagen gel extraction kit (28704). We eluted this DNA in 30µl of molecular grade water heated to 65°C applied to multiple columns. Eurofins manufactured (and salt-free purified) the following oligo which matched one of the subclones: TTGTATAGAAAAGTTGTACTGTGGTATCACTCGTC ACTTGCGTAGAACGCTTGGA AGGCTCAGCTAAGGTGCAATGGTAGGCGTGTACGGTG. We then used Gibson assembly (NEB E2611S) followed by precipitation and resuspension to extract a plasmid construct. We transformed OneShot Stbl3 (Invitrogen C737303) bacterial cells with this construct.

We transfected LentiX to produce lentivirus. We collected 10ml of virus containing Tu2% from four 15cm plates, each day for three days. We passed all Tu2% through a 0.45µm filter (Millipore Sigma SE1M003M00) and then ultracentrifuged it at 39,800rcf for 30min at 4°C. We resuspended virus in 7ml of Tu2%. WM989 subclones were all spinfected with this virus, with 50,000 cells total plated per well in a 6-well plate. We transduced WM989 cells with 1ml of resuspended virus. We centrifuged the 6-well plate at 600rcf for 25min. We left cells in the virus overnight. The next day we replaced the virus with 0.5µg/ml doxycycline (Sigma D9891-25g) containing Tu2%. Two days later we collected cells for FACS analysis using a FACSJazz instrument.

### Development of dabrafenib and trametinib resistant cells

We transduced WM989 A6-G3 with PB-TRE-Cas9-VPR with CloneSweeper virus using spinfection at 600rcf for 25min, with 200,000 cells plated per well in eight 6-well plates. Each well contained 300µl of CloneSweeper virus. We left cells overnight in virus, we then replaced the virus with fresh Tu2% the following morning. A day later we sorted for EGFP positive cells and then plated them into two 10cm plates, with 140,000 cells in each plate. Five days later we moved cells from each plate into one 15cm dish each, two days after that we moved each pool of cells into four 15cm dishes. Finally, five days later, we collected cells from each pool for two semi-independent resistance experiments. This expansion corresponded to ∼8.5 doublings. We centrifuged 4,000,000 cells from each pool into cell pellets for analysis of lineage barcode distribution prior to treatment via genomic DNA sequencing. We froze 10 cryovials of live cells, from each pool, with approximately 4,500,000 cells per vial, for later lineage barcode enrichment. We plated 800,000 cells per plate in two 15cm dishes per pool. A day after plating we replaced Tu2% with Tu2% with 250nM dabrafenib (Cayman CAYM-16989-10) and 2.5nM trametinib (SelleckChem S2673 871700-17-3) added. 42 days later we collected cell pellets for genomic DNA extraction.

### Sequencing of resistant genomic DNA

We extracted genomic DNA from these pellets using a Qiamp DNA Mini Kit (Qiagen 51304). We eluted DNA using two passes of 50µl of nuclease-free water each (Ambion AM9937) heated to 65°C. We added a CloneSweeper calibration “ladder” to cell pellets prior to DNA isolation. This ladder is composed of barcoded subclones of the internally generated G9 subclone of the mouse melanoma line M10M6^40^. This ladder is composed of 10 distinct barcodes with 20 cells each, 4 distinct barcodes with 100 cells each and 4 distinct barcodes with 200 cells each. We amplified lineage barcodes and added Illumina adapters using NEBNext Ultra II Q5 (NEB M0544L). To account for the higher number of cells in pre-treatment pellets we conducted 32 50µl reactions on pre-treatment gDNA while conducting 2 50µl reactions on gDNA extracted from resistant cells. Each reaction contained 500ng of genomic DNA and 500nM of each primer (Table S1, S2). The following PCR cycle was used: Initial denaturation at 98°C for 30 seconds; 25 cycles of: denaturation at 98°C for 10 seconds; annealing and extension at 65°C for 40 seconds and a final extension at 65°C for 5 minutes. We used double-sided magnetic bead (Beckman Coulter A63881) purification with 0.6× and 1× DNA to bead solvent ratios to remove primers and genomic DNA from the PCR reactions. We confirmed appropriate amplicon size and purity using a high-sensitivity bioanalyzer (Agilent 5067-4626). We sequenced this product using an Illumina P1 kit on a NextSeq 1000.

### Analysis of genomic DNA

We extracted CloneSweeper barcodes from fastqs using code we updated for our new lineage barcoding library (https://github.com/SydShafferLab/BarcodeAnalysis_v2_gDNA). We included reads that matched 90% or more of nucleotides immediately downstream of the barcode. We clustered together CloneSweeper barcodes with 3 or less Levenshtein distance from another, we then assigned barcodes to the most frequent raw barcode sequence using Starcode^41^. We identified the mean number of reads for 200 cell ladder rungs, we used this number as a threshold that identified cells primed to resistance that we tested CloneSweeper’s cell enrichment capacity on. Shannon and Simpson indices were calculated using the vegan package in R.

### Generation of enrichment mini-library

Twist Bioscience generated 21 pooled oligos (Table S3) targeting 21 CloneSweeper barcodes. We prepared a 50µl Gibson assembly (NEB E2611S) reaction to add oligos to linearized (via method outlined earlier) VB230528-1280dkp (Figure 1F), with the following: 250ng linearized plasmid and 285.25 fmol pooled oligo. We precipitated the resulting DNA and resuspended it in 10µl water. We applied the same procedure to Gibson assembly without oligos added, as a control. We then transformed Endura electrocompetent bacteria (Bioresearch Technologies 71003-036) with 2µl of concentrated DNA per 25µl bacteria. We estimated 370,000 independent transformations based on colonies plated on ampicillin plates at 1:10,000 for the enrichment mini-library and 1:1000 for the negative control. We isolated plasmids using a maxi prep (Invitrogen K210006) from bacteria grown in 1.2L of LB broth containing 100µg/ml ampicillin. To test the representation of this enrichment mini-library we generated constructs with Illumina adapters by PCR. We used NEBNext Ultra II Q5 (NEB M0544L) with 10pg plasmid and 500nM of each primer (Table S4) with the following: Initial denaturation at 98°C for 30 seconds then 25 cycles of denaturation at 98°C for 10 seconds, annealing at 71°C for 30 seconds, extension at 72°C for 30 seconds, followed by a final extension at 72°C for 2 minutes. We purified these constructs using a Qiagen PCR purification kit (28104) and sequenced on an Element Biosciences Aviti machine.

We plated LentiX cells near 75% confluency in 49 15cm plates. We used one plate for the matched mScarlet3 plasmid generated earlier. We transfected the other 48 plates with enrichment mini-library plasmid. For the one plate we prepared two tubes of 1.25ml of Opti-MEM (Gibco 31985062) with 25µg recovery plasmid, 18.75µg pPax2 plasmid and 12.5µg VSVG plasmid in one tube and 200µl 1mg/ml PEI (Polyscience 23966-1) in the other tube. We added 2.5ml of the combined solution dropwise to the 15cm plate. For the enrichment mini-library plates we added the following to a total 60ml Opti-MEM: 1200µg enrichment mini-library plasmid, 900µg pPax2 plasmid and 600µg VSVG. We added 800µl of 1mg/ml PEI to another total of 60ml Opti-MEM. After combining and incubating for 15min we added 2.5ml of solution per plate dropwise to the enrichment mini-library plates. We replaced DMEM with 12ml of Tu2% approximately 6.5 hours after addition of transfection solution. Then we collected virus containing media every 6-14 hours after. Upon collection we immediately filtered using 0.45µm filters (Fisher Scientific FB12566505) and stored the media at 4°C. After five collections virus we serially ultracentrifuged media at 39,810 rcf for 30min at 4°C. We resuspended virus from the single-barcode targeting plasmid in 2ml Tu2% and enrichment mini-library virus in 20ml Tu2%.

### Enrichment of primed cells and 10x scRNA-seq

We thawed two vials of live-frozen WM989 cells from our resistance experiment and plated them into a 15cm dish. We simultaneously thawed cells from the subclone with a mismatched CloneSweeper barcode. Two days later we plated 200,000 cells per well into 6-well plates: one well was used for the mismatched cells, one for untransduced cells (used as a control for enrichment mini-library independent expansion of lineages) and finally we plated 48 wells for transduction with enrichment mini-library. We added 1.5ml of “matched” plasmid virus to the mismatched well. We added 16ml total of enrichment mini-library virus across 48 wells. We spinfected cells for 25min at 600rcf and left them in virus overnight. The next morning we replaced virus with 0.5µg/ml doxycycline (Sigma D9891-25g) containing Tu2%, in all wells. Two days later we collected cells for FACS Aurora sorting. Only EGFP positive cells were sorted in the control protocol. We centrifuged and resuspended control protocol cells and diluted them in 0.04% BSA (B2518-100mg) containing PBS (Corning 21-031-CV) aiming for 15,000 cells in the 10x protocol. We sorted enrichment protocol cells into EGFP+/mScarlet3- and EGFP+/mScarlet3+ tubes. We used cells with the mismatched CloneSweeper barcode and mScarlet plasmid as a threshold to discriminate mScarlet3 negative and positive cells, only considering cells brighter than all of these cells as mScarlet3 positive. We centrifuged enrichment protocol cells in such a way that all EGFP+/mScarlet3+ (27,800 cells total according to the cytometer, adjusting based on the discrepancy between the cytometer’s count and manually counting in other samples we estimated an actual number of 13,900) were retained while EGFP+/mScarlet3-cells were added such that a total of 25,500 cells were obtained and resuspended in 17µl of 0.04% BSA (B2518-100mg) containing PBS (Corning 21-031-CV). 15,000 cells could be loaded onto the 10x Chromium. We used the Chromium GEM-X Single-Cell 3’ protocol v4 to sequence these pools. We processed enrichment mini-library negative and positive cells in separate lanes. We sequenced fragments twice using an Element Biosciences AVITI with AVITI 2×75 Sequencing Kits Cloudbreak FS High Output (860-00015).

### Analysis of scRNA-seq and identification of CloneSweeper barcodes

We used CellRanger 8.0.1 to extract gene expression data from fastqs. We then loaded expression data into Seurat v5. We removed any cells with 200 or less or 9000 or more RNA features. We only kept cells with more than 10,000 total RNA UMIs. Finally, we filtered out any cells with 10% or more mitochondrial gene UMIs out of total gene UMIs. We updated code to isolate CloneSweeper barcodes directly from 10x scRNA-seq unmapped sequencing data (https://github.com/SydShafferLab/BarcodeAnalysis_v2_10X). We input reads that matched the CloneSweeper sequence either 16bp upstream or 16bp downstream of the lineage barcode into Starcode^41^. We clustered CloneSweeper barcodes with 3 or less Levenshtein distance from another together. We obtained UMIs from post-clustering CloneSweeper barcodes. At this point our analysis contained a total of 17,172 cells. We eliminated cells with three or more UMIs in more than one CloneSweeper barcode from further analysis, reducing our cell number to 15,975. We then removed cells without any CloneSweeper barcodes with three or more UMIs, leading to a further reduction to 13,544 cells. Of the remaining cells, we only retained cells with a difference of 3 or more between the highest UMI count and the second highest UMI count. This led to a final cell count of 12,996 in our analysis. We then log normalized gene expression with a scale factor of 10,000. We then scaled data and determined variable features. We applied a PCA on the resulting data. We ran FindNeighbors with 10 dimensions and FindClusters with a resolution of 0.5 and then applied a 10 dimension UMAP.

### SCENIC analysis of inferred transcription factor activity

We ran SCENIC^42^ on raw UMI counts. We used only thresholded cells from the enrichment protocol. We inferred gene regulatory networks followed by generation of a context file. Finally, we assigned area under the curve values for each resulting transcription factor activity and corresponding cell. To assess the differences between primed and unprimed lineages we used Wilcoxon tests with Bonferroni corrections on the resulting AUCs.

## Data availability

scRNA-seq data will be deposited on GEO

Imaging, flow cytometry and genomic DNA sequencing data will be deposited on Zenodo.

## Code availability

Code will be made available on Github

## Acknowledgements

We thank all members of the Shaffer Lab for their valuable feedback and support. We would like to thank the genomics core at Wistar as well as the flow cytometry core at Children’s Hospital of Pennsylvania for assistance with experiments. S.M.S. acknowledges support from the NIH Director’s Early Independence Award DP5OD028144, the Institute for Regenerative Medicine (IRM) at the University of Pennsylvania from the Larry and Mickey Magid award, and the American Cancer Society (RSG-23-1152597-01-CDP). S.M.S. is the Bakewell Foundation Innovator of the Damon Runyon Cancer Research Foundation (DRR-81-24). S.M.S. acknowledges support from NIH NIGMS (R01GM149671). We would like to thank Benjamin L. Emert for advice with respect to cloning lineage barcode libraries. microtube-open-translucent icon drawn by Servier https://smart.servier.com/ is licensed under CC-BY 3.0 Unported (https://creativecommons.org/licenses/by/3.0/).

---

**Supplementary Figure 1:**
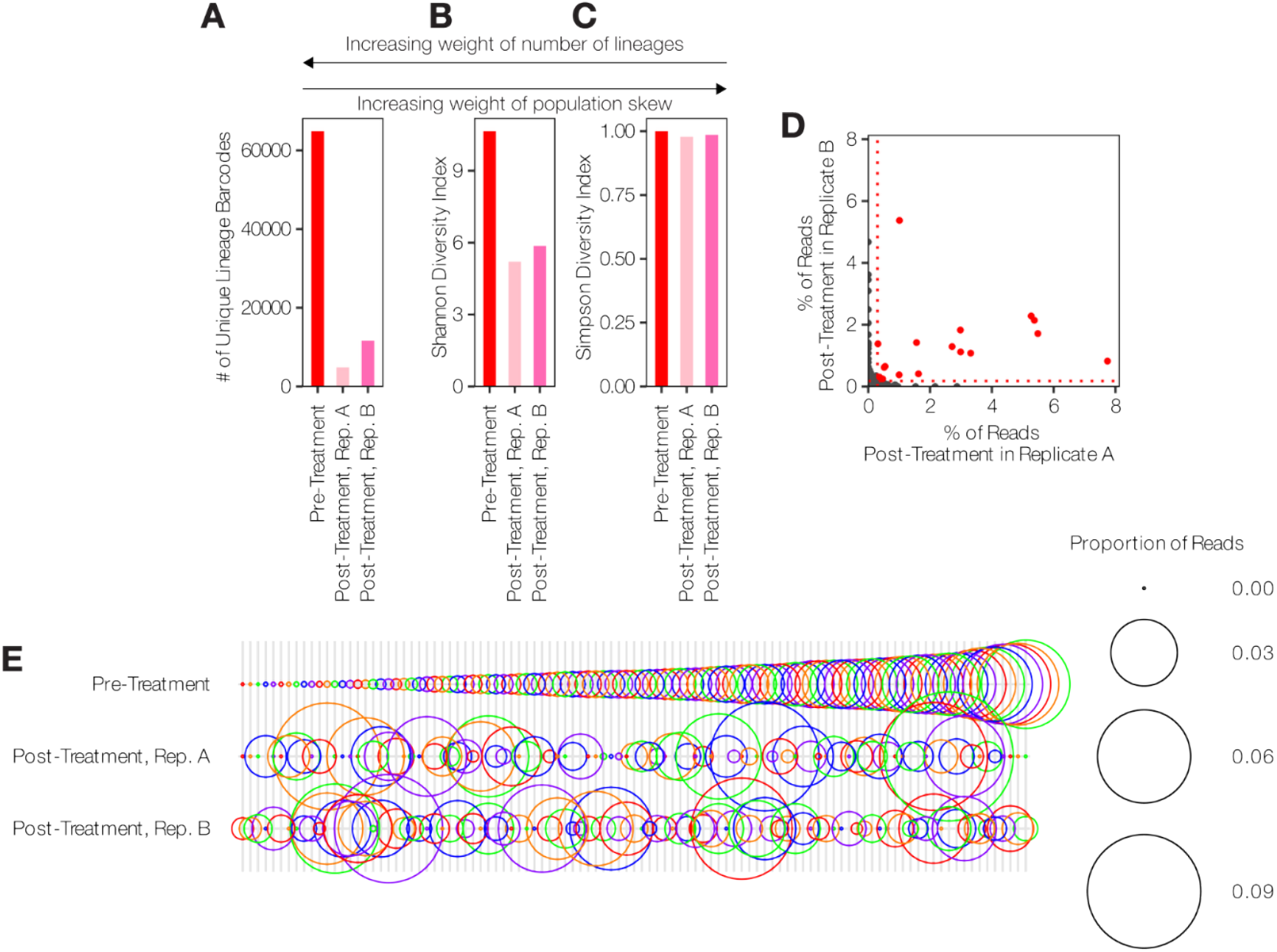
Semi-independent replication of resistance dynamics from Figure 2. **A-C**. Diversity metrics for semi-independent replicate of resistance development tracking CloneTracer clones in dabrafenib and trametinib (see Figure 2). **A**. Number of lineage barcodes detected. **B**. Shannon diversity of lineage barcodes. **C**. Simpson index of lineage barcodes. Metrics in B-C only included lineage barcodes with 2 or more reads in either pre-treatment or post-treatment samples. **D**. Percentage of CloneSweeper barcode reads. The mean percentage of spiked-in cells with 200 cells are plotted as dotted red lines. **E**. Bubble plot showing the proportion of reads of a lineage barcode out of all lineages passing the 200 cell threshold in at least one sample. Only lineage barcodes passing this threshold are shown. Colors cycle across adjacent lineage barcodes for ease of identification.

**Supplementary Figure 2:**
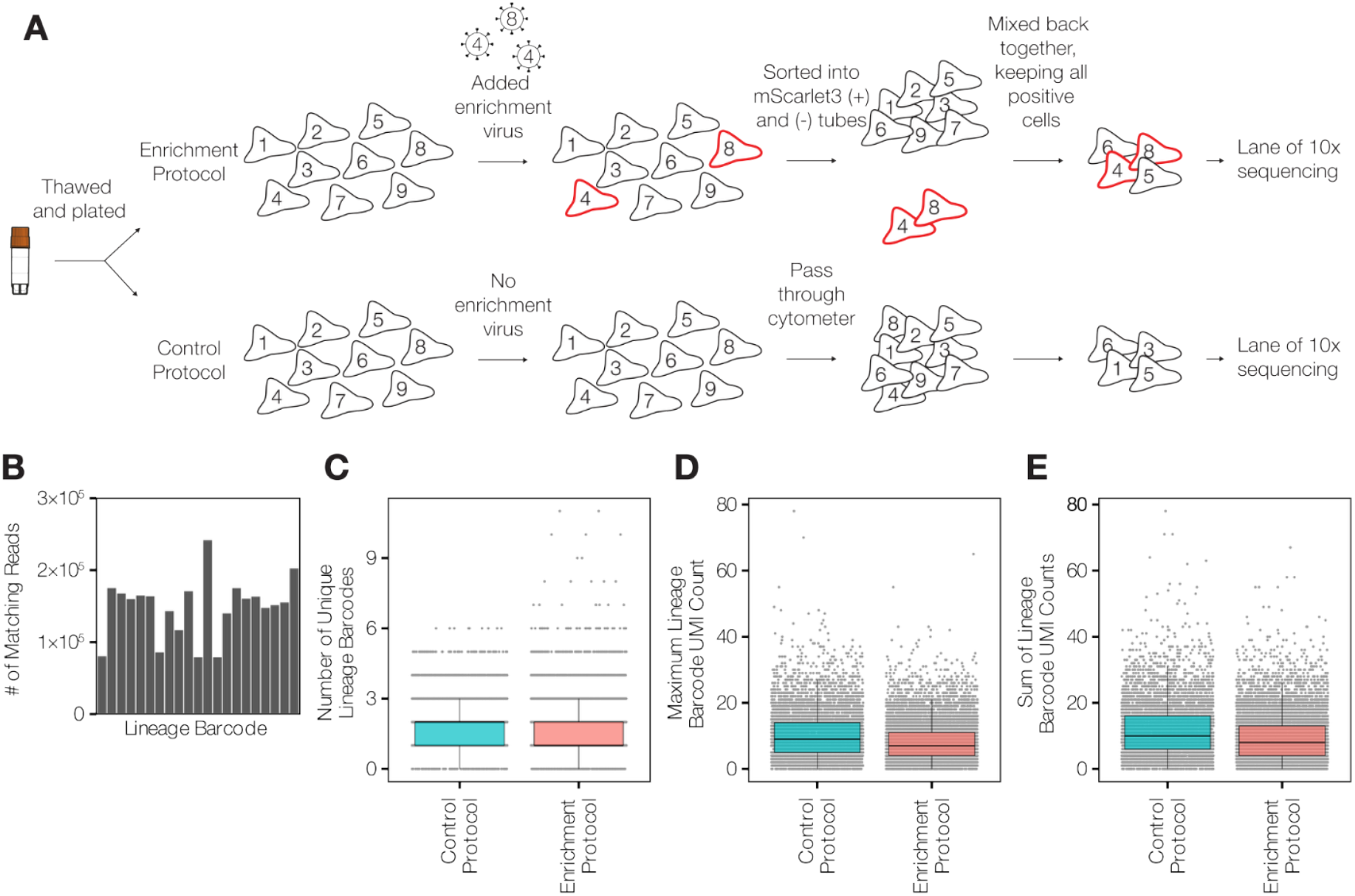
Enrichment workflow design and scRNA-seq quality control. **A**. Detailed schematic of the cell sorting strategy for scRNA-seq. To ensure sufficient material for 10x Genomics scRNA-seq loading, sorted mScarlet3+ (enriched) cells were spiked with mScarlet3-cells to reach a target loading density. **B**. Next-generation sequencing of the plasmid “mini-library” prior to transduction, confirming the presence of all 21 targeted lineage barcodes (barcodes are shown in the same order as in Figure 3B). **C-E**. Lineage barcode metrics from scRNA-seq. **C**. Number of unique lineage barcodes per cell (≥1 UMIs). **D**. Maximum UMI counts across lineage barcodes in each cell. **E**. Total UMI count across all lineage barcodes in each cell. **C-E**. Boxplots show median and interquartile ranges (IQR), whiskers show 1.5× the IQR.

**Supplementary Figure 3:**
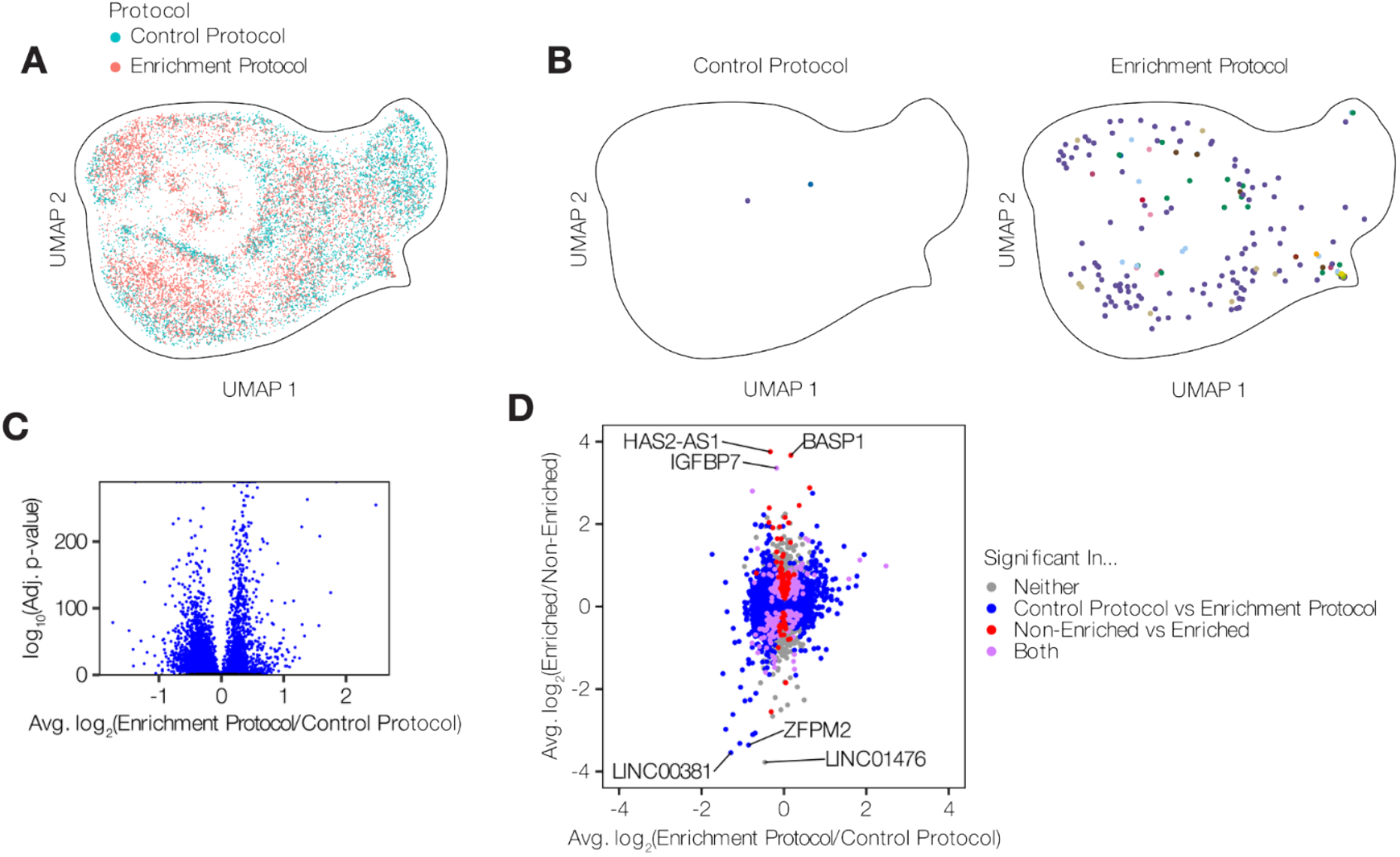
Enrichment does not introduce significant transcriptional bias. **A**. UMAP projection of scRNA-seq data (without cell cycle regression) comparing the cells from the control protocol (blue) to those from the enrichment protocol (pink). **B**. The same UMAP highlighting the location of cells carrying the 21 targeted lineage barcodes from the control versus enrichment protocols. Outline is aligned to correspond exactly to the outline in A. **C**. Volcano plot describing differences in gene expression between cells from the control and enrichment protocols. We calculated p-values based on Wilcoxon tests with Bonferroni corrections. Only genes detected (UMIs ≥ 1) in 5% of cells in both groups are shown. Cells with enriched lineage barcodes were removed in this analysis for consistency. **D**. Plot showing average log_2_ fold change in expression, simultaneously showing comparisons from Figure 4A and Figure S6C, we considered p-value < 0.05 significant. Only genes detected (UMIs ≥ 1) in 5% of cells in both groups from either comparison are shown.

**Supplementary Table 1:**
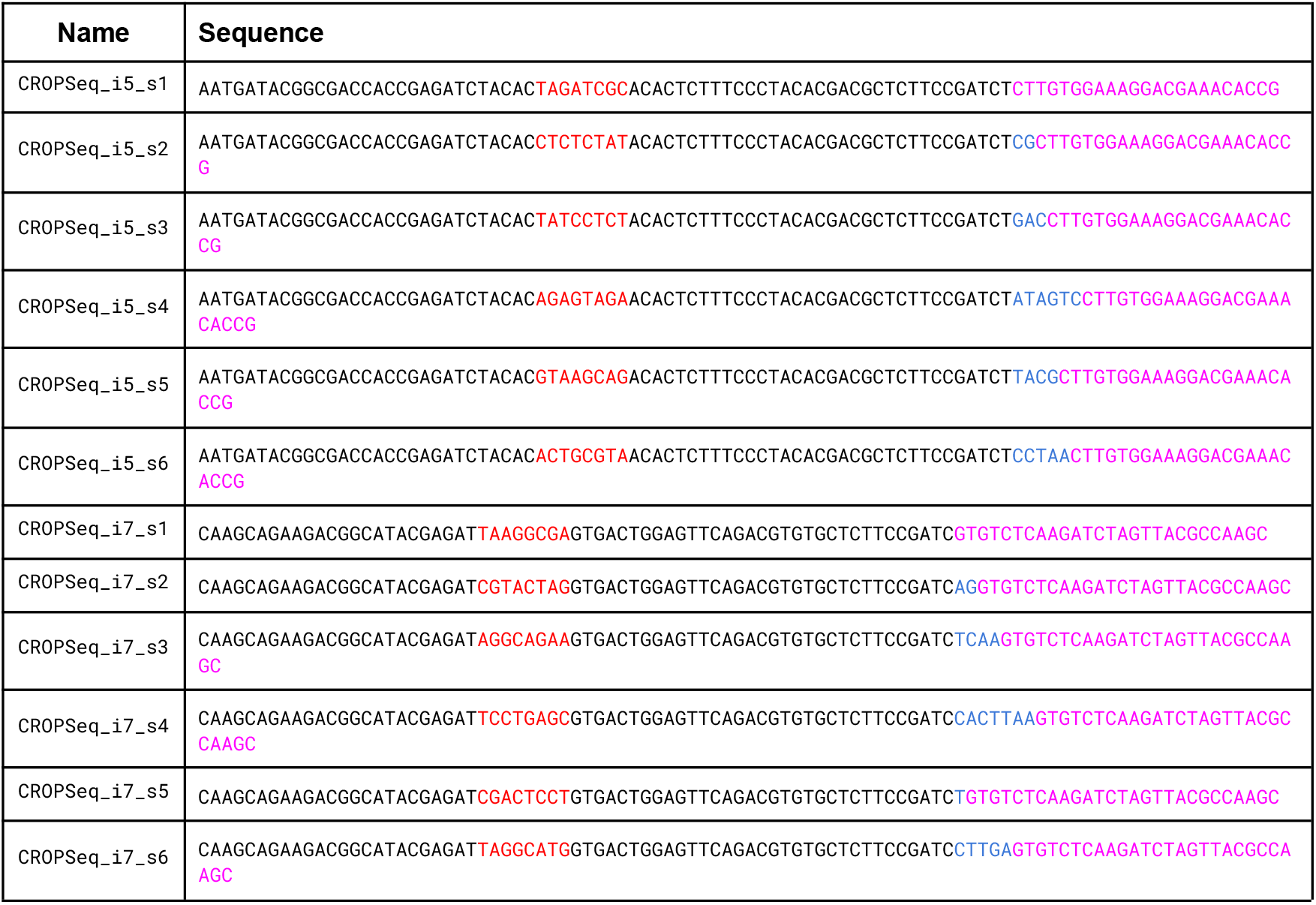
Primer sequences used for amplification of CloneSweeper barcodes from gDNA.

**Supplementary Table 2:** Specific primers used for amplification of gDNA.

| Sample | i5 primer | i7 primer |
| --- | --- | --- |
| Pre-Treatment (Main Figures) | s4 | s1 |
| Post-Treatment A (Main Figures) | s5 | s1 |
| Post-Treatment B (Main Figures) | s6 | s2 |
| Pre-Treatment (Supplemental Figures) | s3 | s1 |
| Post-Treatment A (Supplemental Figures) | s1 | s5 |
| Post-Treatment B (Supplemental Figures) | s2 | s6 |

**Supplementary Table 3:**
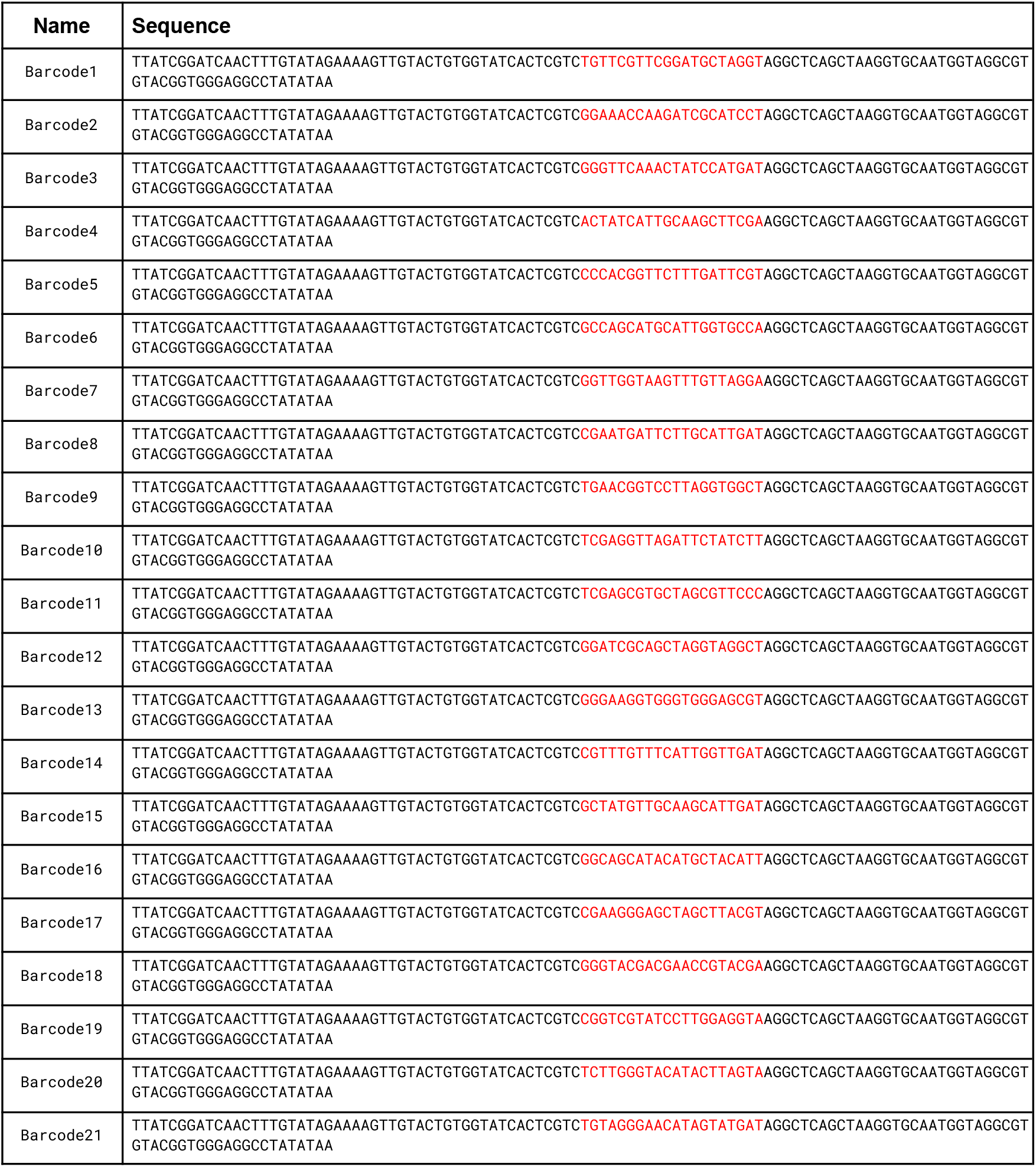
Sequences of pooled oligos for cloning of enrichment library.

**Supplementary Table 4:**
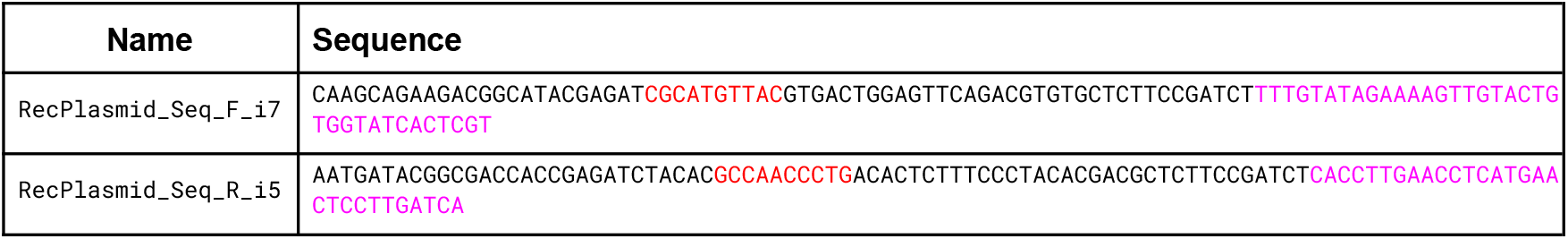
Primers for sequencing enrichment library.

